# Challenging the loss of complexity theory: insights from the neuromuscular system in chronic obstructive pulmonary disease

**DOI:** 10.1101/2025.10.10.681623

**Authors:** Cyril Chatain, Sofiane Ramdani, Nicolas Paleiron, Fanny Cucchietti Waltz, Asmaa Jobic, Jean-Marc Vallier, Mathieu Gruet

**Affiliations:** Inserm UMR1093 – Cognition Action & Plasticité Sensorimotrice (CAPS), Université Bourgogne Europe, UFR des Sciences du Sport, Dijon, France; Laboratoire d’Informatique, de Robotique et de Microélectronique de Montpellier (LIRMM), Université de Montpellier, Centre National de la Recherche Scientifique (CNRS), Montpellier, France; Laboratoire Jeunesse - Activité Physique et Sportive - Santé (J-AP2S), Université de Toulon, Toulon, France; Service de Pneumologie, Hôpital d’Instruction des Armées Saint-Anne, Toulon, France; Délégation à la Recherche Clinique et à l’Innovation (DRCI), Centre Hospitalier Intercommunal de Toulon – La Seyne sur Mer (CHITS), Toulon, France

**Keywords:** muscle functioning, dual-task, COPD, cognitive control, neuromuscular fatigability, interference

## Abstract

**Background:** According to the “loss of complexity” theory, aging and disease are expected to reduce complexity of physiological outputs, thereby limiting the system’s adaptability. However, it remains unclear whether this concept applies to the neuromuscular system in people with chronic obstructive pulmonary disease (pwCOPD). This study aimed to challenge the loss of complexity hypothesis by assessing the regularity, as well as the steadiness and the accuracy, of force production during submaximal isometric contractions in pwCOPD compared to healthy individuals.

**Methods:** Seventeen pwCOPD and seventeen age- and sex-matched healthy participants performed submaximal isometric contractions of the knee extensors at six target forces, ranging from 10 to 60% of their maximal voluntary contraction (MVC). Regularity of force signals was assessed using sample entropy (SampEn) and percentage of determinism (DET) from the recurrence quantification analysis. Steadiness and accuracy were quantified using the coefficient of variation (CV) and the root-mean-square error (RMSE), respectively.

**Results:** PwCOPD exhibited 26.5% lower MVC than healthy individuals. Despite this muscular weakness, no significant main effect of group or interaction effect (group × contraction intensity) was observed for SampEn, DET, CV and RMSE, suggesting a preserved force control in pwCOPD at all assessed force levels.

**Conclusion:** Our results indicate that the loss of complexity theory may not apply in moderate COPD, at least for the neuromuscular system. These findings suggest that neuromuscular alteration associated with COPD may not be sufficient to impair the complexity of force output, questioning the universality of the loss of complexity theory.

## 1. Introduction

The loss of “complexity” theory, first proposed by (Lipsitz & Goldberger, 1992), suggests that aging and disease can lead to a reduction in the number of components within physiological systems and in the richness of their interactions (Sleimen-Malkoun et al., 2014). This reduction in internal complexity is thought to compromise the system’s adaptative capacity, i.e., its ability to respond quickly and effectively to internal and/or external perturbations, ultimately bringing individuals closer to threshold of frailty (Lipsitz, 2002; Vaillancourt & Newell, 2003).

Such changes can be estimated by analyzing the fluctuations of physiological signals generated by the system of interest. In particular, examining the temporal structure of fluctuations using nonlinear time series analysis, also known as “measures of complexity” (e.g., entropy measures, analyses of recurring patterns), can provide valuable insights into the functional status and the underlying dynamics of the system (Costa et al., 2005; Goldberger et al., 2002; Manor & Lipsitz, 2013).

Applied to neuromuscular system, this conceptual framework suggests that loss of force “complexity”, expressed by a reduction in the irregularity or unpredictability of torque or force signals, reflects a breakdown in the interaction between neural and muscular components involved in force production (Sleimen-Malkoun et al., 2014). Numerous studies showed that aging (Fiogbé et al., 2021; Vaillancourt & Newell, 2003) and neurological disorders such as Parkinson’s disease or stroke (Chow & Stokic, 2014; Vaillancourt et al., 2001) are associated with reduced complexity of force or torque output during isometric muscle contractions. These findings support the idea that aging and pathological conditions may drive the neuromuscular system toward more stereotyped and less adaptative behavior.

In this context, chronic obstructive pulmonary disease (COPD) represents a compelling model to further explore this theory. Although primarily characterized by airflow limitation, it is well documented that COPD, through primary (i.e., disease related) and secondary (i.e., related to treatment or comorbidities) factors, is also associated with various systemic manifestations. Among them, neuromuscular impairments stand as a hallmark of the disease. At peripheral level, structural changes include a decreased proportion and size of type I muscle fibers, reduced oxidative capacity and lower mitochondrial density (Maltais et al., 2014). At central level, accumulating evidence suggests that people with COPD (pwCOPD) also exhibit impaired neuromuscular function due to reduced cortical activity, excitability and voluntary activation (Alexandre et al., 2014, 2020). Altogether, these impairments can compromise the integrity of neuromuscular control processes, which may manifest as an increased variability (i.e., greater deviations in force magnitude around the target) and/or reduced complexity (i.e., more regular and less adaptable temporal structure of force fluctuations) in force output.

Alexandre et al., (2014) reported that pwCOPD exhibit impaired submaximal force control compared to age- and sex-matched healthy individuals during knee extensors contractions performed at 10, 30 and 50% of maximal voluntary force (MVF). However, the authors assessed force control using only an inaccuracy metric (i.e., root-mean-square error, quantifying average error from the target), which provides limited information about the temporal structure of force fluctuations. As emphasized by Pethick et al., (2021a), a more comprehensive approach to evaluating force control should incorporate both magnitude-based indices (e.g., coefficient of variation, root-mean square error) and complexity measures (e.g., entropy, recurrence analysis), in order to capture the multifaceted nature of neuromuscular function.

Beyond challenging the loss of complexity theory in the context of a multisystem disease, demonstrating impairments in neuromuscular complexity in pwCOPD could contribute to the emergence of new indicators of physiological adaptability. Such markers may be particularly relevant and sensitive for discriminating between populations or evaluating the effectiveness of interventions acting across different physiological systems (e.g., exercise-based rehabilitation). For instance, Ramdani et al., (2013) reported that measures from recurrence quantification analysis applied to center of pressure time series could successfully differentiate older fallers from non-fallers. Similarly, Carnavale et al., (2020) showed that entropy measures derived from force signals during submaximal contractions were able to distinguish frail from non-frail older adults, and that approximate entropy could even discriminate pre-frail from frail individuals. In addition, it has been demonstrated that multiscale entropy applied to postural sway would be more sensitive than conventional center of pressure metrics to detect the effects of Tai Chi training program in healthy older adults (Wayne et al., 2014).

Therefore, the aim of this study was to investigate the force control of the knee extensors in pwCOPD compared to healthy individuals across a range of submaximal intensities (i.e., from 10 to 60% MVF). To this end, we used a comprehensive set of indices: coefficient of variation (CV), root-mean-square error (RMSE), Sample Entropy (SampEn), and Recurrence Quantification Analysis (RQA). Based on the loss of complexity framework and findings provided by Alexandre et al., (2014), we hypothesized that pwCOPD would exhibit less complex and more variable force output. Identifying such impairments could help guide the development of targeted rehabilitation strategies aimed at improving the quality and adaptability of neuromuscular control.

## 2. Methods

### 2.1 Participants and experimental design

Seventeen pwCOPD and seventeen age- and sex-matched healthy control participants were enrolled in this study. The main inclusion criteria were: (1) forced expiratory volume in the first second < 80% of predicted values (for pwCOPD only); (2) stable state (i.e., without exacerbations) for more than 15 days (for pwCOPD only); (3) without oral or systemic corticosteroids (> 0.5 mg.kg^-1^.day^-1^ for more than 7 days) treatment (for pwCOPD only); (4) without oxygen therapy (for pwCOPD only); (5) body mass index < 30 kg.m^-2^ (for all participants); (6) without uncorrected severe vision or hearing impairments (for all participants); (7) without vigorous physical activity with a frequency of more than three sessions per week (for all participants).

The study received ethical approval from an ethic committee (CPP Ile de France III, 2019-A01986-51) and registered at www.clinicaltrials.gov (NCT06871670). All participants provided written informed consent prior to their participation in the study and were free to withdraw at any time.

Data presented in this study were collected from a larger project (NCT04028973) aiming to investigate the influence of cognitive load on muscle endurance and neurophysiological adjustments during a fatiguing task between pwCOPD and healthy individuals (Chatain et al., 2024). In the current study, only some data (i.e., force signals) recorded during the first of three visits were retained for analysis. These data have not been previously explored and offer a distinct perspective from the initial project. It should be stressed that all procedures completed during the first visit (e.g., questionnaire completion, muscle contractions) were performed using the same sequence and same timing for each volunteer. All participants were asked to refrain from caffeine and vigorous or prolonged physical activity 24h prior to each visit.

### 2.2 Dynamometry and data acquisition

During the first visit, participants were familiarized with the equipment and study procedures. Participants were seated in an adjustable custom-built chair with hips and knees at 130° and 90° of flexion, respectively. The lower leg was attached to a force sensor (F2712 200 daN, Celians MEIRI, France) approximately 5 cm above the malleoli of the ankle joint with an inextensible padded Velcro strap, while a belt secured firmly across the waist to prevent any extraneous movements during isometric contractions. Force signals were recorded using BIOPAC MP150 (BIOPAC Systems, Inc., Santa Barbara, CA, USA) and sampled at 2 kHz with AcqKnowledge software (version 4.1, BIOPAC Systems, Inc., Santa Barbara, CA, USA).

### 2.3 Experimental procedures

After installation and familiarization with isometric contractions, participants performed a standardized warm-up of the knee extensors involving 3 min intermittent isometric contractions at increasing force levels. After a 2 min recovery period, volunteers were asked to perform at least three maximal voluntary contractions (MVCs) of approximately 5 s duration with 1-min rest intervals (Figure 1). If a gradual increase was observed (i.e., increase >5% between the third MVC and the largest of the first two), additional MVCs were performed to obtain the true MVF. Strong verbal encouragements were given during each MVC to ensure maximal effort.

**Figure 1.**
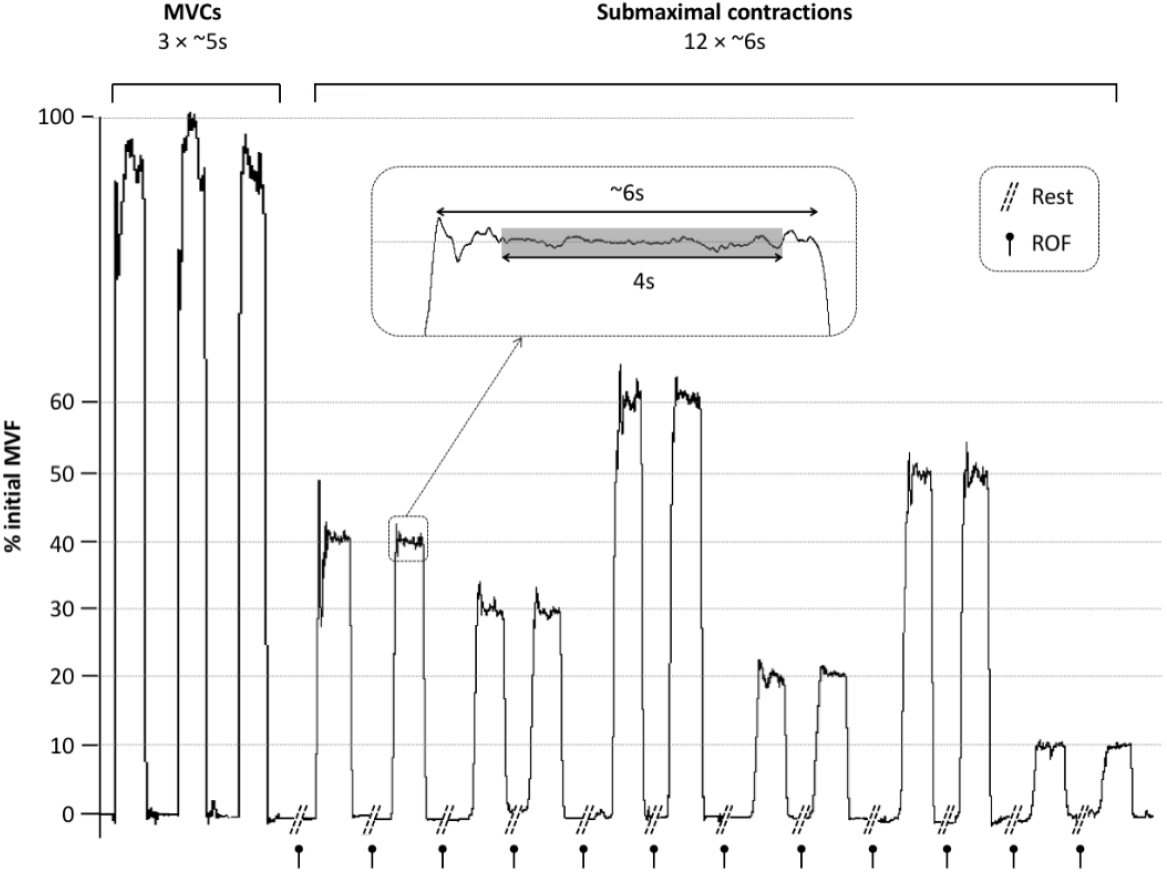
Example of force signal recorded during the force control section of the protocol including the accomplishment of several MVCs and the submaximal contractions performed at different intensities. Gray area shows the window retained for signal analysis for each submaximal contraction. MVC: Maximal voluntary contraction; MVF: Maximal voluntary force; ROF: Rating of fatigue scale.

Then, after 2 min of rest, participants were asked to perform two submaximal isometric knee extensions at different intensity levels, i.e., 10%, 20%, 30%, 40%, 50% and 60% MVF (Figure 1). Submaximal force levels were calculated based on the best MVC previously performed. The force signal feedback was displayed on a TV screen placed in front of the participants (∼ 150 cm distance) with a resolution of 3.2% MVC.cm^-1^. During each submaximal contraction, participants were asked to maintain the produced force as accurately and steadily as possible during ∼ 6s. For a given intensity (e.g., 10% MVF), both contractions were performed in a row but several intensities were accomplished in a random order. The recovery time between each submaximal contraction was not standardized in terms of duration, but participants were allowed to perform a contraction only if their rate of fatigue scale (ROF) score was ≤ 2/10 (0 corresponding to *not fatigued at all*, a score between 2 and 3 corresponding to *a little fatigued* and a score of 10 corresponding to *total fatigue and exhaustion*) (Micklewright et al., 2017). This procedure was preferred over a standardized recovery time (e.g., 30s between each contraction) to limit the effect of potential differences in muscle recovery between participants and populations. Nevertheless, participants were instructed that recovery durations typically range between 15-30s for low force levels, and 30s or more for higher contractions intensities. The experimenter also monitored the pace and provided guidance when necessary to ensure relative consistent testing conditions between participants.

### 2.4 Data analysis

For each submaximal contraction, the 4 s steadiest and closest to the target force level by visual inspection were extracted using AcqKnowledge software (Figure 1). Then, signal processing was performed with MATLAB software (version R2023b, MathWorks, Natick, MA, USA) by using codes from Goldberger et al., (2000) and the Cross Recurrence Plot Toolbox (version 5.22) developed by Marwan, (2004). Force signals were off-line filtered using 6^th^ order zero-phase low-pass digital Butterworth filter with cut-off frequency of 20 Hz and down-sampled at 500 Hz.

The coefficient of variation (CV) and the root-mean-square error (RMSE) from the target were computed to quantify force variability and force accuracy, respectively. CV (%) was calculated as the ratio between standard deviation (SD) and mean force using the formula (1) while RMSE (%) was calculated as the absolute difference between the average produced force and the target force expressed as a percentage of the target using the formula (2):

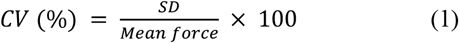

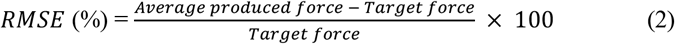

We made the choice to select relative measures of force variability and accuracy rather than absolute indicators (i.e., variability expressed in SD and accuracy in N·m) because they are preferred to compare groups which present different strengths capacities.

Force signal regularity was quantified using two approaches: the sample entropy (SampEn) and the percentage of determinism (DET) from recurrence quantification analysis (RQA). Prior to regularity analyses, the very low frequency trend was removed for each signal using Empirical Mode Decomposition (EMD) method (Huang et al., 1998). In short, EMD is a non-linear and data driven method allowing to decompose a signal into different components with locally defined frequency bands. EMD can also be used to limit first-order nonstationarity of a signal by removing the lowest frequency component (also called residual) [see Flandrin et al. (2004) for details and Chatain et al. (2021) for example of application on force signals]. This procedure was performed to ensure that potential differences of regularity between contraction levels and groups are not mainly explained by very low-frequency trends. Prior to SampEn and RQA computations, signals were standardized (i.e., zero mean and unit standard deviation).

A detailed description of SampEn computation (formula 3) is available in Richman & Moorman (2000) and Lake et al., (2002). In short, it measures the regularity of time series by quantifying the negative natural logarithm of the conditional probability that a sequence, also called dimension *d*, of similar points remains similar with an additional point (i.e., *d*+1) according to a tolerance *r* (so-called radius). In formula (3), B represents the number of similar sequences in dimension d and A represents the number of sequences that remain similar in dimension d+1 within a time series of length *N*. In the current study, the embedding dimension *d* was set to 3 for both SampEn and RQA (see below) based on results from the global false nearest neighbor algorithm (Chatain et al., 2021; Kennel et al., 1992). In agreement with previous studies (Chatain et al., 2020; Pethick et al., 2015) and recommendations from Forrest et al., (2014) tolerance *r* was set to 0.1 for SampEn computation.

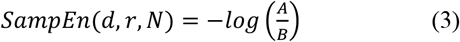

RQA method (Marwan et al., 2007; Marwan & Kraemer, 2023) is based on the reconstruction of the phase space from a time series produced by a dynamical system (i.e., force signal from neuromuscular system in the current study) following Taken’s delay embedding theorem (Takens, 1981) using a dimension *d*, a delay *τ* and the following formula (4):

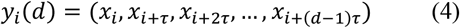

where *x*_*i*_ is the *i*^th^ point of the original temporal series, *y*_*i*_*(d)* the *i*^th^ sequence (also called time delay vector) of dimension *d*.

Then RQA consists of reconstructing a recurrence matrix, also called recurrence plot (RP) which corresponds to a 2-dimensional binary representation of the recurrence states of the underlying dynamics using formula (5):

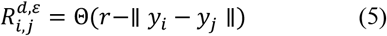

where Θ is the Heaviside function, ∥ ∥ is the Euclidean norm and *y*_*i*_ and *y*_*j*_ represent the state vectors of the system at two instants *i* and *j*, respectively. In short, RP is composed of black and white points. A black point (i.e., a recurrence point) is placed at coordinates (*i,j*) when the distance between the state vectors *y*_*i*_ and *y*_*j*_ is below the threshold or tolerance *ε*, meaning that the state of the system at time *i* is closed to its state at time *j*. Otherwise, no recurrence point is plotted.

Finally, DET quantifies the proportion of recurrence points within the RP that form diagonal lines of at least length *l*_*min*_. These diagonals indicate that similar states recur over a period of time, reflecting a degree of predictability or regularity of time series. DET is computed using equation (6) where *P(l)* is the histogram of diagonal lines of length *l*.

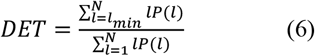

Several empirical rules have been proposed to select the tolerance *r* for the reconstruction of RP (e.g., to choose *ε* value leading to a fixed recurrence point density within the RP) (Marwan et al., 2007). In the current study, we used a more recent approach proposed by Medrano et al., (2021) based on a statistical method using a Kernel Density Estimator to optimize the tolerance selection. In contrast, there is no rigorous method allowing to objectively determine the *l*_*min*_ parameter. Therefore, in order to prevent a threshold effect for the estimation of DET (i.e., to avoid reaching the maximal value of 100%) *l*_*min*_ was set to 50. Delay *τ* used to reconstruct the phase space was set to 20 according to previous work (Chatain et al., 2021).

It is of note that SampEn and DET computations were performed by slightly varying the input parameters (i.e., *d, r, L*_*min*_, *τ*) to confirm the consistency of the results.

### 2.5 Statistical analysis

All statistical procedures were performed on SPSS (version 27.0, IBM Corp., Armonk, NY, USA). For all variables, the normality assumption was inspected both analytically and visually using Shapiro-Wilk and Q-Q plot, respectively. Levene’s test was used to assess the assumption of homogeneity of variances. Participants’ characteristics and CV, RMSE, SampEn and DET (when compared for specific intensity) were compared between groups using independent sample t-test when parametric assumptions were met (i.e., normality and homogeneity of variances). When normality was met but homogeneity of variances was violated, Welch’s test was used. When normality assumption was not met, the non-parametric Mann-Whitney test was applied. Effect sizes were obtained using the measure of Cohen’s *d* (interpreted as follow: 0.2 small; 0.5 moderate; 0.8 large) and rank-biserial correlation coefficient for parametric and non-parametric analysis, respectively (Cohen, 1992; Tomczak & Tomczak, 2014).

To detect effects of group, intensity or interaction (group x intensity), CV, RMSE, SampEn and DET were compared using linear mixed models (LMMs) with one within-subject factor (intensity) and one between subject factor (group). The effect of contraction was included in the models as a repeated measure but was not included as a fixed factor as it was not of interest in the analyses. Therefore, the fixed factors of the LMMs were intensity, group and interaction with a random effect structured by subjects. If a main effect or interaction was detected, LMMs were followed by pairwise comparison tests using the Bonferroni correction.

As previously mentioned, in addition to LMMs, group comparisons at each specific intensity were also performed. These analyses were performed: i) because the emphasis was on the group differences more than the effect of intensity or interaction and ii) because it allowed the identification of potential differences for specific intensities that might attenuated or not fully captured by interaction in the LMMs. This approach was also in line with some studies that focused exploration of force fluctuations at only one intensity (e.g., Heffernan et al., 2009; Pancera et al., 2022; Pereira et al., 2024). For these analyses, the values obtained during both contractions at each intensity for different metrics were averaged to obtain a single value per subject and per intensity.

All data are presented as mean ± SD within the text and figures. Mean differences (MD) and 95% confidence intervals (95% CI) were also computed. For all analyses, statistical inferences were based on a *p*-value threshold of 0.05.

## 3. Results

### 3.1 Participants characteristics

Main characteristics of the participants are presented in Table 1. No difference was reported for the different variables, except for FEV_1_ and MVF which were significantly lower in pwCOPD than in control participants.

**Table 1.** Main characteristics of participants. BMI: Body mass index; FEV_1_: Forced expiratory volume during the first second; MVF: Maximal voluntary force; MMSE: Mini mental state examination; pwCOPD: People with chronic obstructive pulmonary disease. *(n=16). $: statistical analysis performed using non-parametric approach.

|  | *pwCOPD* | *controls* | *p-value* | *Effect size* |
| --- | --- | --- | --- | --- |
| Gender M/F | 10/7 | 10/7 |  |  |
| Age (years) | 69.9±7.0 | 68.8±6.6 | 0.617 | 0.173 |
| Weight (kg) | 68.7±9.9 | 72.1±15.6 | 0.453 | 0.261 |
| Height (cm) | 166.9±7.6 | 171.4±9.4 | 0.136 | 0.524 |
| BMI (kg.m<sup>-2</sup>) | 24.7±3.5 | 24.4±4.3 | 0.826 | 0.076 |
| FEV<sub>1</sub> (L)<sup>\$</sup> | 1.5±0.5 | 3.3±0.8* | <0.001 | 2.620 |
| FEV<sub>1</sub> (% predicted) | 59.3±12.2 | 121.8±13.4* | <0.001 | 4.890 |
| MVF (N.m) | 141.5±52.0 | 187.7±63.0 | 0.030 | 0.799 |
| MMSE (0-30)<sup>\$</sup> | 28.6±1.0 | 28.9±1.2 | 0.405 | 0.166 |

### 3.2 Mean force

Prior to data analysis, we confirmed that all participants respected task instructions and that there were no significant differences between groups in terms of relative mean force production (all *p*>0.15; Figure 2).

**Figure 2.**
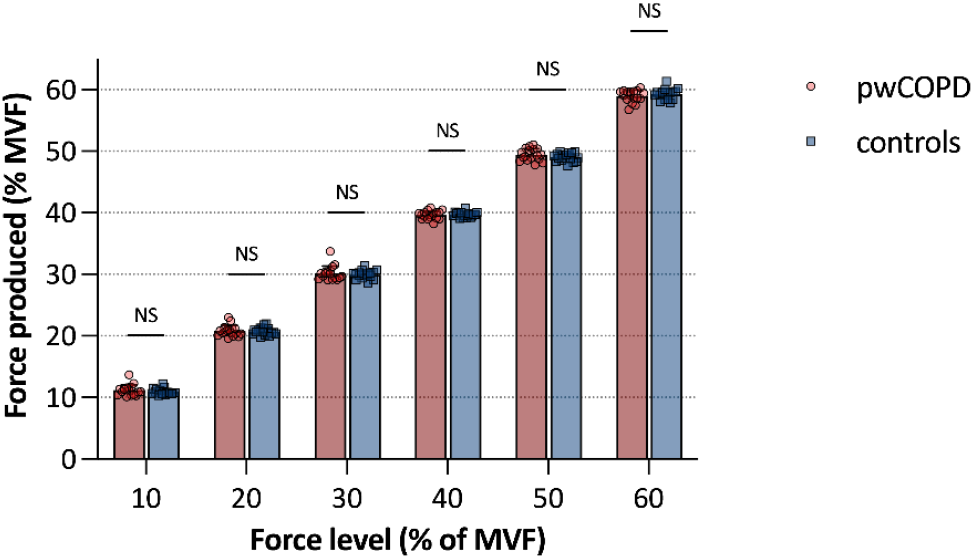
Mean relative force produced as a function of force intensities for pwCOPD (in red) and healthy participants (in blue). Each point represents individual data. Error bars represent the standard deviation. NS : non-significant.

### 3.3 Steadiness and accuracy of force production

For CV, LMM revealed a significant effect of force level [F(5,194.8)=13.1; *p*<0.001] without main effect of group [MD=0.31; 95%CI=[-0.08;0.70]; F(1,32.0)=2.7; *p*=0.11] or interaction effect [F(5,194.8)=0.7; *p*=0.65] (Figure 3). CV was significantly higher at 10% force level compared to all other force levels (all *p*<0.005).

**Figure 3.**
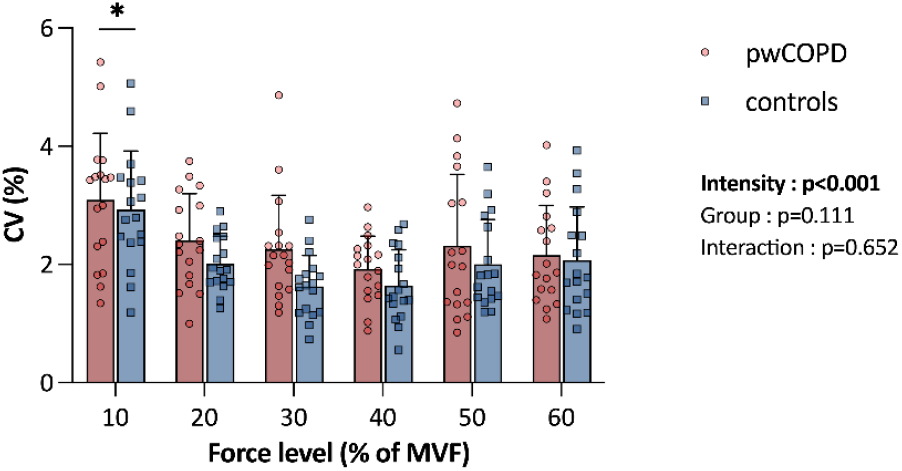
Effect of force intensity on coefficient of variation (CV) for both people with COPD (pwCOPD) and controls. *: significantly higher than all other force levels.

LMM revealed a significant main effect of force level for RMSE [F(5,273.7)=24.0; *p*<0.001] without main effect of group [MD=0.83; 95%CI=[-0.71;2.37]; F(1,41.5)=1.2; *p*=0.28] or interaction effect [F(5,273.7)=1.2; *p*=0.33] (Figure 4). RMSE was significantly higher at 10% force level compared to all other force levels (all *p*<0.001). RMSE was also significantly different at 20% force level compared to all other force levels (all *p*<0.05).

**Figure 4.**
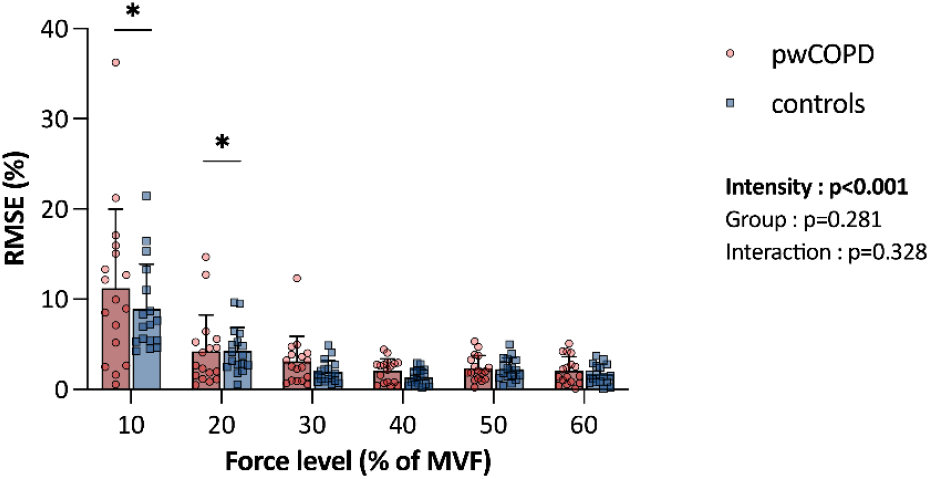
Effect of force level on root mean square error (RMSE) for both people with COPD (pwCOPD) and controls. *: significantly different to all other force levels.

When comparisons between groups were performed separately for each force intensity, only one statistically significant difference was reported for CV at 30% force level (*p*=0.020; Table 2). No significant difference between groups was reported for RMSE (all *p*>0.06; Table 2).

**Table 2.**
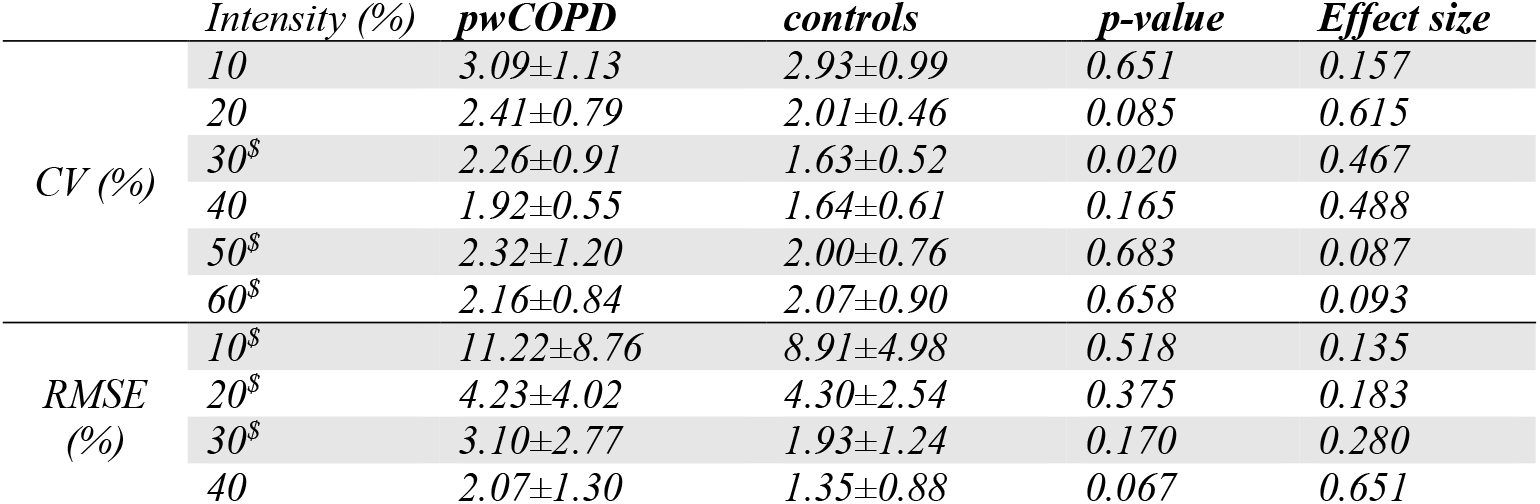
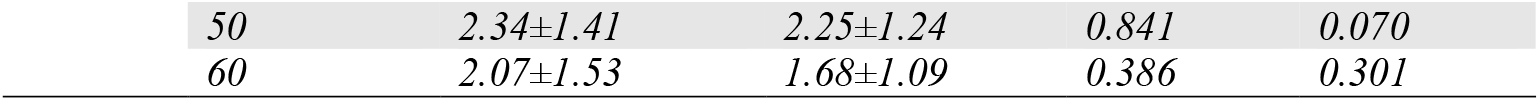
Independent analysis of differences between pwCOPD and controls groups for CV and RMSE at each force level. $: statistical analysis performed using non-parametric approach.

### 3.4 Regularity of force production

LMM revealed a significant main effect of force level for SampEn [F(5,200.2)=5.1; *p*<0.001] without main effect of group [MD=0.009; 95%CI=[-0.021;0.040]; F(1,32.0)=0.4; *p*=0.53] or interaction effect [F(5,200.2)=1.4; *p*=0.24] (Figure 5). SampEn was significantly lower at 10% and 20% force levels compared to 40%, 50% and 60% force levels (all *p*<0.05).

**Figure 5.**
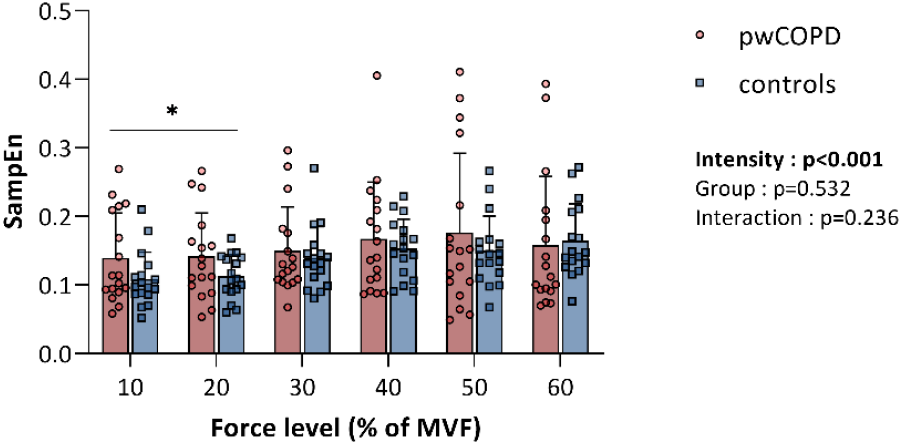
Effect of force level on sample entropy (SampEn) for both people with COPD (pwCOPD) and controls. *: significantly different to 40%, 50% and 60% force levels.

LMM revealed a significant main effect of force level for DET [F(5,203.1)=4.4; *p*<0.001] without main effect of group [MD=0.031; 95%CI=[-0.115;0.053]; F(1,32.0)=0.6; *p*=0.46] or interaction effect [F(5,203.1)=1.3; *p*=0.27] (Figure 6). DET was significantly higher at 10% force level compared to 40% force level (*p*<0.05). DET was also significantly higher at 20% force level compared to 40%, 50% and 60% force levels (all *p*<0.05).

**Figure 6.**
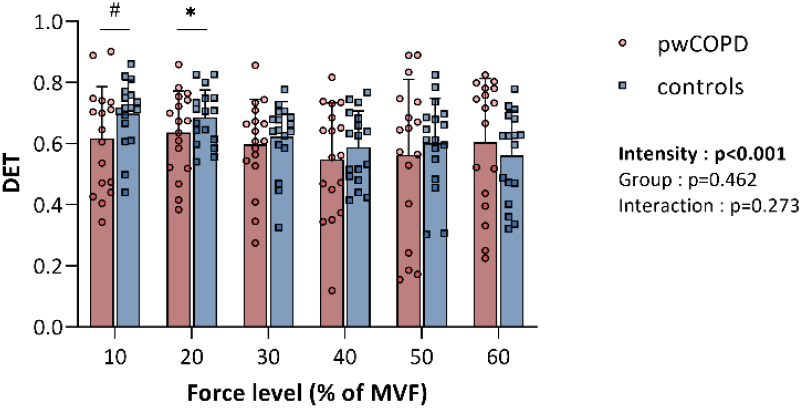
Effect of force level on determinism (DET) for both people with COPD (pwCOPD) and controls. ^#^: significantly different to 40% force level; *: significantly different to 40%, 50% and 60% force levels.

When comparisons between groups were performed separately for each force intensity, no significant difference between groups was reported for SampEn and DET (all *p*>0.10) (Table 3).

**Table 3.** Independent analysis of differences between pwCOPD and controls for SampEn and DET at each force level. $: statistical analysis performed using non-parametric approach.

|  |  | ***pwCOPD (n=17)*** | ***controls (n=17)*** | ***p-value*** | ***Effect size*** |
| --- | --- | --- | --- | --- | --- |
| *SampEn* | *10<sup>\$</sup>* | *0.14±0.06* | *0.12±0.04* | *0.182* | *0.273* |
|  | *20* | *0.15±0.05* | *0.12±0.03* | *0.144* | *0.514* |
| | *30<sup>\$</sup>* | *0.15±0.05* | *0.15±0.04* | *0.999* | *0.235* |
|  | *40* | *0.17±0.06* | *0.16±0.04* | *0.679* | *0.144* |
|  | *50* | *0.17±0.09* | *0.16±0.05* | *0.651* | *0.157* |
|  | *60* | *0.15±0.07* | *0.17±0.04* | *0.486* | *0.242* |
| *DET* | *10* | *0.62±0.17* | *0.70±0.11* | *0.105* | *0.575* |
|  | *20* | *0.64±0.14* | *0.69±0.09* | *0.216* | *0.433* |
| | *30<sup>\$</sup>* | *0.60±0.15* | *0.62±0.11* | *0.518* | *0.135* |
|  | *40* | *0.55±0.18* | *0.59±0.12* | *0.461* | *0.256* |
| | *50<sup>\$</sup>* | *0.56±0.25* | *0.60±0.15* | *0.838* | *0.045* |
| | *60<sup>\$</sup>* | *0.60±0.21* | *0.56±0.15* | *0.205* | *0.260* |

## 4. Discussion

This study aimed to assess force control during submaximal isometric contractions of the knee extensors performed at different intensities in pwCOPD, compared to age- and sex-matched healthy participants, using a set of complementary indices that capture steadiness, accuracy and complexity of force output. In line with the “loss of complexity” theory, we hypothesized that pwCOPD would display less complex and more variable force output compared to healthy participants. Contrary to our assumption, our results showed no changes in the temporal structure of force signals induced by COPD suggesting that the loss of complexity theory is not as straightforward as conventionally admitted, at least, for the neuromuscular system. Coupled with the results obtained with measures of magnitude variability, it appears that COPD does not significantly impaired force control.

### 4.1 Effect of COPD on force control production

As a preliminary observation, pwCOPD exhibited 26.5% lower muscle force than healthy controls. This is consistent with previous force deficit reported in the literature in moderate pwCOPD which typically ranges from 20% to 30% (Alexandre et al., 2014; Mador & Bozkanat, 2001; Maltais et al., 2014). Regardless of whether this difference in maximal force production originates from peripheral and/or central components of neuromuscular function, it confirms that we assessed different neuromuscular systems. Nevertheless, the use of relative intensities during submaximal contractions guaranteed that force conditions were similar between pwCOPD and healthy participants. In this respect, we found that participants in both groups were equally able to perform the submaximal contractions according to the instructions (i.e., no differences between groups in terms of relative mean force production).

The absence of a main group effect on all force control indices (i.e., steadiness, accuracy, complexity) suggests that the global force control production is not impaired in pwCOPD. However, when compared at specific intensities, pwCOPD showed significantly higher CV than healthy participants at 30% MVF, suggesting that a loss of force steadiness induced by the disease may emerge under particular task constraints. Interestingly, values of CV obtained in the current study in pwCOPD (i.e., approximately 2%) were substantially lower that those reported by Pancera et al., (2022). Indeed, using a similar task (i.e., submaximal isometric contractions of the knee extensors performed at 30% MVF), the authors found CV around 11%. Although the absence of a control group in their study limits the interpretation, these elevated values suggest that force steadiness can be remarkably impaired in some patients, a phenomenon that was not observed in the present study. The relatively low CV values observed here in pwCOPD may partially explain the limited differences found between groups at other contraction intensities and for other indices of force control. It is worth noting that patients in our study presented similar FEV_1_ compared to those involved in the study of Pancera et al., (2022), suggesting that force steadiness (and more widely the force control production) impairments may not be solely explained by disease severity.

The absence of decrease in force production accuracy in pwCOPD (i.e., absence of significant differences between groups for RMSE) even when compared at specific intensities, contrasts with previous results reported by Alexandre et al., (2014). Indeed, the authors showed a higher force inaccuracy (i.e., higher RMSE) in pwCOPD during knee extensors contractions performed at 10, 30 and 50% MVF. The absence of statistical difference reported here may, at least in part, be explained by the important interindividual variability, especially in pwCOPD group at low force intensities.

Regarding complexity measures (i.e., SampEn and DET), no significant group differences were found across intensities. This finding does not support the “loss of complexity” theory which postulates that aging and disease leads to less complex dynamics of physiological signals thereby reducing the adaptation abilities of a system to external perturbations (Goldberger et al., 2002; Lipsitz, 2004; Lipsitz & Goldberger, 1992). While different studies reported reduced force or torque complexity in several diseases (e.g., stroke, Down syndrome, Parkinson’s disease) (Chow & Stokic, 2014; Heffernan et al., 2009; Vaillancourt et al., 2001), our findings suggest that this may not extend to moderate COPD. Although the present study is focused on the effect of COPD, our results are in line with findings of Vieluf et al., (2015) obtained in healthy elderly individuals. Indeed, the authors reported no statistical differences in force complexity between healthy young and older adults during flexion of the index finger using a derivative metrics of SampEn, namely Multi-Scale Entropy. Importantly, in the current study, our two distinct non-linear measures of complexity (i.e., SampEn and DET from RQA), which capture different aspects of temporal structure, converge toward the same conclusion. This strengthens the interpretation that a systematic loss of complexity may not be as evident or universal as often assumed, particularly in the context of certain pathological conditions such as moderate COPD.

One possible explanation to the absence of loss of complexity in pwCOPD could be linked to the nature and the extent of neuromuscular impairments. In contrast to neurological disorders such as Parkinson’s disease or stroke in which clear structural and functional alterations of central neural system exist, these impairments in moderate COPD may be more subtle. Although the inflammation and circulating cytokines in COPD could be implicated in neural damages and increased risk of neurodegenerative disorders (e.g., Alzheimer’s or Parkinson’s disease) (Liao et al., 2015), the importance of central nervous system impairments in moderate stages of the disease might not be sufficient to significantly change the complexity of force output. Future studies could explore this assumption by comparing patients with COPD alone to pwCOPD and a co-existent neurodegenerative disorder.

Assuming that severe COPD are at increased risks of brain impairments (Borson et al., 2008), one may expect increased likeliness of loss of neuromuscular complexity compared to mild-to moderate disease profile. In addition, compensatory neuromuscular strategies developed by some patients could have contributed to preserve complexity during submaximal tasks.

Moreover, it is of note that these findings were observed by confronting pwCOPD and healthy participants to similar constraints, by adapting the disease-related strength deficit through the use of relative force. This approach is in line with previous studies assessing difference in force complexity between different populations (e.g., young *vs*. old adults or healthy *vs*. pathological individuals) (Heffernan et al., 2009; Ofori et al., 2018; Vaillancourt et al., 2001; Vaillancourt & Newell, 2003; Vieluf et al., 2015) and seems appropriate to isolate neuromuscular control from the confounding effect of muscle weakness. Indeed, to compare individuals based on absolute force levels would result in disproportionate neuromuscular demands for weaker individuals such pwCOPD. Therefore, it can be assumed that if the task difficulty was scaled in absolute force level, differences in complexity between populations could have emerged. Exploring this assumption in future work could help to distinguish the effects of age- or disease-related muscle weakness from those related to force complexity as such.

### 4.2 Effect of contraction intensity on force control production

As expected, our results showed a significant effect of contraction intensity on all force control indices. Specifically, relative force variability (i.e., CV) significantly decreased with increasing contraction intensity, with a higher variability observed at 10% MVF compared to all other intensities. This result is consistent with previous literature showing a marked reduction in CV from low to moderate intensities followed by a stabilization from moderate to high intensities, in healthy individuals (Christou et al., 2002; Ofori et al., 2018). To the best of our knowledge, our results extend for the first time this observation to pwCOPD. In terms of force accuracy, a significant decrease in RMSE was reported with increasing intensity, aligning with previous finding in both healthy (Henry et al., 2025) and pwCOPD (Alexandre et al., 2014). This effect of contraction intensity on force variability and accuracy could be explained, at least in part, by the resolution of the visual feedback which, when expressed in absolute terms, was lower at low contraction intensities (Pereira et al., 2024). Therefore, participants may have more difficulty to correct small deviations in force. In addition to visual feedback constraints, it has been shown that some physiological mechanisms are also involved in the decrease of relative force variability such as the reduction in motor unit discharge rate variability (Moritz et al., 2005), the changes in slow common oscillations in the discharge times (Negro et al., 2009) and synchronization of motor units (Taylor et al., 2003).

Regarding the complexity of force output, assessed by two regularity metrics (i.e., SampEn and DET from RQA), our results showed a significant increase in irregularity of force production with increasing force levels. These results are in partial agreement with previous studies, although there is no consensus about the effect of the intensity of force production on force output complexity. For instance, Fiogbé et al., (2021) also reported an increase in SampEn values of knee extensors force output from 15% to 40% MVF in healthy young and old adults. In contrast, Pethick et al., (2021b) showed a decrease of the same metric from 25% to 100% MVF and Ofori et al., (2018) found no changes of Approximate Entropy (a measure close to SampEn) across intensities from 2% to 95% MVF during isometric contractions of knee extensors in young adults. Such discrepancies between studies remain unclear. It is worth noting that in the present study, both conceptually distinct indicators of regularity (i.e., SampEn and DET) revealed similar trends, thereby strengthening the robustness of our findings. Future studies are needed to better understand how contraction intensity modulates force output complexity. This may help determine the most effective intensity range to promote the interaction of neuromuscular system components and maximize the benefits of exercise training.

Finally, although the mechanisms behind the force output complexity are poorly understood, it was suggested that an optimal complexity could be explained by the simultaneous availability of two neural strategies for force production (i.e., recruitment of additional motor units and the modulation of their discharge rate) allowing the neuromuscular system benefits from maximal flexibility to adjust the force output (Slifkin & Newell, 1999). Our findings suggest that, at least until 60% MVF, this flexibility continues to increase during knee extensors contractions.

### 4.3 Limitations and perspectives

One limitation of the present study is that most participants had moderate COPD. The relatively small sample size did not allow for stratification based on disease severity. However, this sample remains consistent with previous studies investigating force complexity in both healthy and clinical populations (e.g., Ofori et al., 2018; Vaillancourt et al., 2001; Vieluf et al., 2015). While the current study was not specifically powered to test hypotheses regarding the impact of COPD severity on force complexity, the relative homogeneity of our sample with respect to disease severity and the consistency across two distinct nonlinear approaches, both pointing toward preserved complexity, strengthens that the absence of significant group difference in unlikely due to insufficient statistical power. Nonetheless, future research should investigate whether disease severity is associated with progressive impairments in force control production.

Another limitation of the present study is that it implies only force signals recordings during muscle contraction, limiting the interpretation of the underlying mechanisms that may contribute to changes in force complexity with increasing contraction intensity. In this context, future studies could benefit from using techniques such as high-density electromyography to better understand the neuromuscular mechanisms underlying the force complexity output, particularly regarding motor unit activity. The use of computational models allowing to simulate force signals could also represent a valuable approach to further elucidate the mechanisms involved.

Finally, a recent meta-analysis highlighted a relationship between force steadiness and functional capacities in older adults (Camacho-Villa et al., 2025). Therefore, it would be of interest to investigate whether impairments in force control are linked to functional limitations in pwCOPD. The approach used in the present study, combining several force control indices (i.e., steadiness, accuracy and complexity) may provide a useful framework for such investigation.

## 5. Conclusion

To the best of our knowledge, this is the first study to investigate force control production in pwCOPD using a comprehensive set of indices, including both non-linear measures and traditional linear outcomes. Overall, our findings suggest that force control is not markedly impaired in pwCOPD compared to healthy individuals. Surprisingly, the absence of effect on complexity measures suggests a similar organization of the system underlying force control between healthy participants and pwCOPD. Future studies are needed to better understand the generalization of the loss of complexity theory to neuromuscular function in a multisystem disease such as COPD, extending our investigations to more severe disease profile and to COPD with co-existent neurodegenerative disorders.

## Abbreviations

BMI: Body mass index
COPD: Chronic obstructive pulmonary disease
CV: Coefficient of variation
DET: Percentage of determinism
EMD: Empirical mode decomposition
FEV_1_: Forced expiratory volume in the first second
LMM: Linear mixed models
MD: Mean difference
MMSE: Mini-mental state examination
MVC: Maximal voluntary contraction
MVF: Maximal voluntary force
PwCOPD: People with chronic obstructive pulmonary disease
RMSE: Root-mean-square error
ROF: Rating-of-fatigue scale
RP: Reccurence plot
RQA: Recurrence quantification analysis
SampEn: Sample entropy
SD: Standard deviation

## References

Alexandre, F., Heraud, N., Oliver, N., & Varray, A. (2014). Cortical Implication in Lower Voluntary Muscle Force Production in Non-Hypoxemic COPD Patients. PLoS ONE, 9(6), e100961. 10.1371/journal.pone.0100961

Alexandre, F., Héraud, N., Tremey, E., Oliver, N., Bourgouin, D., & Varray, A. (2020). Specific motor cortex hypoexcitability and hypoactivation in COPD patients with peripheral muscle weakness. BMC Pulmonary Medicine, 20(1). 10.1186/S12890-019-1042-0

Borson, S., Scanlan, J., Friedman, S., Zuhr, E., Fields, J., Aylward, E., Mahurin, R., Richards, T., Anzai, Y., Yukawa, M., & Yeh, S. (2008). Modeling the impact of COPD on the brain. International Journal of Chronic Obstructive Pulmonary Disease, 3(3), 429–434. 10.2147/COPD.S2066

Camacho-Villa, M. A., Giráldez-García, M. A., Sevilla-Sanchez, M., Rivera-Mejía, S. L., & Carballeira, E. (2025). Relationship Between Force Steadiness and Functionality in Older Adults: A Systematic Review With Meta-Analysis. Scandinavian Journal of Medicine & Science in Sports, 35(4), e70040. 10.1111/SMS.70040

Carnavale, B. F., Fiogbé, E., Farche, A. C. S., Catai, A. M., Porta, A., & Takahashi, A. C. de M. (2020). Complexity of knee extensor torque in patients with frailty syndrome: a cross-sectional study. Brazilian Journal of Physical Therapy. 10.1016/j.bjpt.2018.12.004

Chatain, C., Gruet, M., Vallier, J.-M., & Ramdani, S. (2020). Effects of nonstationarity on muscle force signals regularity during a fatiguing motor task. IEEE Transactions on Neural Systems and Rehabilitation Engineering, 28(1), 228–237. 10.1109/tnsre.2019.2955808

Chatain, C., Ramdani, S., Vallier, J. M., & Gruet, M. (2021). Recurrence quantification analysis of force signals to assess neuromuscular fatigue in men and women. Biomedical Signal Processing and Control, 68, 102593. 10.1016/J.BSPC.2021.102593

Chatain, C., Vallier, J.-M., Paleiron, N., Cucchietti Waltz, F., Ramdani, S., & Gruet, M. (2024). Muscle endurance, neuromuscular fatigability, and cognitive control during prolonged dual-task in people with chronic obstructive pulmonary disease: a case-control study. European Journal of Applied Physiology. 10.1007/S00421-024-05608-X

Chow, J. W., & Stokic, D. S. (2014). Variability, frequency composition, and complexity of submaximal isometric knee extension force from subacute to chronic stroke. Neuroscience, 273, 189–198. 10.1016/j.neuroscience.2014.05.018

Christou, E. A., Grossman, M., & Carlton, L. G. (2002). Modeling Variability of Force During Isometric Contractions of the Quadriceps Femoris. Journal of Motor Behavior, 34(1), 67– 81. 10.1080/00222890209601932

Cohen, J. (1992). A power primer. Psychological Bulletin, 112(1), 155–159. 10.1037//0033-2909.112.1.155

Costa, M., Goldberger, A. L., & Peng, C.-K. (2005). Multiscale entropy analysis of biological signals. Physical Review E, 71(2), 021906. 10.1103/PhysRevE.71.021906

Fiogbé, E., Vassimon-Barroso, V., Catai, A. M., De Melo, R. C., Quitério, R. J., Porta, A., & Takahashi, A. C. D. M. (2021). Complexity of Knee Extensor Torque: Effect of Aging and Contraction Intensity. Journal of Strength and Conditioning Research, 35(4), 1050–1057. 10.1519/JSC.0000000000002888

Flandrin, P., Gonçalvès, P., & Rilling, G. (2004). Detrending and denoising with empirical mode decompositions. 2004 12th European Signal Processing Conference, 1581–1584.

Forrest, S. M., Challis, J. H., & Winter, S. L. (2014). The effect of signal acquisition and processing choices on ApEn values: towards a “gold standard” for distinguishing effort levels from isometric force records. Medical Engineering & Physics, 36(6), 676–683.10.1016/J.MEDENGPHY.2014.02.017

Goldberger, A. L., Amaral, L. A., Glass, L., Hausdorff, J. M., Ivanov, P. C., Mark, R. G., Mietus, J. E., Moody, G. B., Peng, C. K., & Stanley, H. E. (2000). PhysioBank, PhysioToolkit, and PhysioNet: components of a new research resource for complex physiologic signals. Circulation, 101(23), E215–E220. http://www.ncbi.nlm.nih.gov/pubmed/10851218

Goldberger, A. L., Amaral, L. A. N., Hausdorff, J. M., Ivanov, P. Ch., Peng, C.-K., & Stanley, H. E. (2002). Fractal dynamics in physiology: alterations with disease and aging. Proceedings of the National Academy of Sciences, 99(Supplement 1), 2466–2472. 10.1073/pnas.012579499

Heffernan, K. S., Sosnoff, J. J., Ofori, E., Jae, S. Y., Baynard, T., Collier, S. R., Goulopoulou, S., Figueroa, A., Woods, J. A., Pitetti, K. H., & Fernhall, B. (2009). Complexity of force output during static exercise in individuals with Down syndrome. Journal of Applied Physiology, 106(4), 1227–1233. 10.1152/JAPPLPHYSIOL.90555.2008,

Henry, M., Darendeli, A., Tvrdy, T., Daneshgar, S., & Enoka, R. M. (2025). Influence of age and feedback modality on the proprioceptive sense of force: insights from motor unit recordings. Journal of Neurophysiology, 133(4), 1103–1115. 10.1152/JN.00486.2024/ASSET/IMAGES/LARGE/JN.00486.2024_F004.JPEG

Huang, N. E., Shen, Z., Long, S. R., Wu, M. C., Shih, H. H., Zheng, Q., Yen, N.-C., Tung, C. C., & Liu, H. H. (1998). The empirical mode decomposition and the Hilbert spectrum for nonlinear and non-stationary time series analysis. Proceedings of the Royal Society of London. Series A: Mathematical, Physical and Engineering Sciences, 454(1971), 903– 995. 10.1098/rspa.1998.0193

Kennel, M. B., Brown, R., & Abarbanel, H. D. I. (1992). Determining embedding dimension for phase-space reconstruction using a geometrical construction. Physical Review A, 45(6), 3403–3411. 10.1103/PhysRevA.45.3403

Lake, D. E., Richman, J. S., Griffin, M. P., & Moorman, J. R. (2002).Sample entropy analysis of neonatal heart rate variability. American Journal of Physiology-Regulatory, Integrative and Comparative Physiology, 283(3), R789–R797. 10.1152/ajpregu.00069.2002

Liao, K. M., Ho, C. H., Ko, S. C., & Li, C. Y. (2015). Increased Risk of Dementia in Patients With Chronic Obstructive Pulmonary Disease. Medicine, 94(23), e930. 10.1097/MD.0000000000000930

Lipsitz, L. A. (2002). Dynamics of stability: the physiologic basis of functional health and frailty. The Journals of Gerontology. Series A, Biological Sciences and Medical Sciences, 57(3), B115–B125. http://www.ncbi.nlm.nih.gov/pubmed/11867648

Lipsitz, L. A. (2004). Physiological complexity, aging, and the path to frailty. Science of Aging Knowledge Environment : SAGE KE, 2004(16). 10.1126/SAGEKE.2004.16.PE16

Lipsitz, L. A., & Goldberger, A. L. (1992). Loss of “complexity” and aging. Potential applications of fractals and chaos theory to senescence. The Journal of the American Medical Association, 267(13), 1806–1809.

Mador, M. J., & Bozkanat, E. (2001). Skeletal muscle dysfunction in chronic obstructive pulmonary disease. Respiratory Research, 2(4), 216–224. 10.1186/RR60/METRICS

Maltais, F., Decramer, M., Casaburi, R., Barreiro, E., Burelle, Y., Debigaré, R., Dekhuijzen, P. N. R., Franssen, F., Gayan-Ramirez, G., Gea, J., Gosker, H. R., Gosselink, R., Hayot, M., Hussain, S. N. A., Janssens, W., Polkey, M. I., Roca, J., Saey, D., Schols, A. M. W. J., … ATS/ERS Ad Hoc Committee on Limb Muscle Dysfunction in COPD. (2014). An official American Thoracic Society/European Respiratory Society statement: update on limb muscle dysfunction in chronic obstructive pulmonary disease. American Journal of Respiratory and Critical Care Medicine, 189(9), e15–e62. 10.1164/rccm.201402-0373ST

Manor, B., & Lipsitz, L. A. (2013). Physiologic complexity and aging: Implications for physical function and rehabilitation. Progress in Neuro-Psychopharmacology and Biological Psychiatry, 45, 287–293. 10.1016/j.pnpbp.2012.08.020

Marwan, N. (2004). Cross Recurrence Plot Toolbox for MATLAB®, Ver. 5.22 (R32.4).https://tocsy.pik-potsdam.de/CRPtoolbox/

Marwan, N., Carmen Romano, M., Thiel, M., & Kurths, J. (2007). Recurrence plots for the analysis of complex systems. Physics Reports, 438(5–6), 237–329. 10.1016/J.PHYSREP.2006.11.001

Marwan, N., & Kraemer, K. H. (2023). Trends in recurrence analysis of dynamical systems. The European Physical Journal Special Topics 2023 232:1, 232(1), 5–27. 10.1140/EPJS/S11734-022-00739-8

Medrano, J., Kheddar, A., Lesne, A., & Ramdani, S. (2021). Radius selection using kernel density estimation for the computation of nonlinear measures. Chaos, 31(8). 10.1063/5.0055797/1077538

Micklewright, D., St Clair Gibson, A., Gladwell, V., & Al Salman, A. (2017). Development and validity of the rating-of-fatigue scale. Sports Medicine, 47(11), 2375–2393. 10.1007/s40279-017-0711-5

Moritz, C. T., Barry, B. K., Pascoe, M. A., & Enoka, R. M. (2005). Discharge rate variability influences the variation in force fluctuations across the working range of a hand muscle. Journal of Neurophysiology, 93(5), 2449–2459. 10.1152/JN.01122.2004

Negro, F., Holobar, A., & Farina, D. (2009). Fluctuations in isometric muscle force can be described by one linear projection of low-frequency components of motor unit discharge rates. The Journal of Physiology, 587(Pt 24), 5925–5938. 10.1113/JPHYSIOL.2009.178509

Ofori, E., Shim, J., & Sosnoff, J. J. (2018). The influence of lower leg configurations on muscle force variability. Journal of Biomechanics, 71, 111–118. 10.1016/j.jbiomech.2018.01.032

Pancera, S., Bianchi, L. N. C., Porta, R., Villafañe, J. H., Buraschi, R., & Lopomo, N. F. (2022). Muscle function and functional performance after pulmonary rehabilitation in patients with chronic obstructive pulmonary disease: a prospective observational study.Scientific Reports, 12(1). 10.1038/S41598-022-20746-Y

Pereira, H. M., Keenan, K. G., & Hunter, S. K. (2024). Influence of visual feedback and cognitive challenge on the age-related changes in force steadiness. Experimental Brain Research, 242(6), 1411–1419. 10.1007/S00221-024-06831-W/FIGURES/2

Pethick, J., Winter, S. L., & Burnley, M. (2015). Fatigue reduces the complexity of knee extensor torque fluctuations during maximal and submaximal intermittent isometric contractions in man. The Journal of Physiology, 593(8), 2085–2096. 10.1113/jphysiol.2015.284380

Pethick, J., Winter, S. L., & Burnley, M. (2021a). Did you know? Using entropy and fractal geometry to quantify fluctuations in physiological outputs. Acta Physiologica, 233(4), e13670. 10.1111/APHA.13670

Pethick, J., Winter, S. L., & Burnley, M. (2021b). Fatigue-induced changes in knee-extensor torque complexity and muscle metabolic rate are dependent on joint angle. European Journal of Applied Physiology, 121(11), 3117–3131. 10.1007/S00421-021-04779-1

Ramdani, S., Tallon, G., Bernard, P. L., & Blain, H. (2013). Recurrence quantification analysis of human postural fluctuations in older fallers and non-fallers. Annals of Biomedical Engineering, 41(8), 1713–1725. 10.1007/s10439-013-0790-x

Richman, J. S., & Moorman, J. R. (2000). Physiological time-series analysis using approximate entropy and sample entropy. American Journal of Physiology-Heart and Circulatory Physiology, 278(6), H2039–H2049. 10.1152/ajpheart.2000.278.6.H2039

Sleimen-Malkoun, R., Temprado, J.-J., & Hong, S. L. (2014). Aging induced loss of complexity and dedifferentiation: consequences for coordination dynamics within and between brain, muscular and behavioral levels. Frontiers in Aging Neuroscience, 6, 140. 10.3389/fnagi.2014.00140

Slifkin, A. B., & Newell, K. M. (1999). Noise, information transmission, and force variability. Journal of Experimental Psychology. Human Perception and Performance, 25(3), 837– 851. 10.1037//0096-1523.25.3.837

Takens, F. (1981). Detecting strange attractors in turbulence. In Dynamical Systems and Turbulence, Warwick 1980, Lecture Notes in Mathematics. ISBN 978-3-540-11171-9. Springer-Verlag. (Vol. 898, pp. 366–381). 10.1007/BFb0091924

Taylor, A. M., Christou, E. A., & Enoka, R. M. (2003). Multiple features of motor-unit activity influence force fluctuations during isometric contractions. Journal of Neurophysiology, 90(2), 1350–1361. 10.1152/JN.00056.2003/ASSET/IMAGES/LARGE/9K0833244007.JPEG

Tomczak, E., & Tomczak, M. (2014). The need to report effect size estimates revisited. An overview of some recommended measures of effect size. TRENDS in Sport Sciences, 21(1), 19–25. https://tss.awf.poznan.pl/The-need-to-report-effect-size-estimates-revisited-An-overview-of-some-recommended,188960,0,2.html

Vaillancourt, D. E., & Newell, K. M. (2003). Aging and the time and frequency structure of force output variability. Journal of Applied Physiology, 94(3), 903–912. 10.1152/japplphysiol.00166.2002

Vaillancourt, D. E., Slifkin, A. B., & Newell, K. M. (2001). Regularity of force tremor in Parkinson’s disease. Clinical Neurophysiology : Official Journal of the International Federation of Clinical Neurophysiology, 112(9), 1594–1603. 10.1016/S1388-2457(01)00593-4

Vieluf, S., Temprado, J. J., Berton, E., Jirsa, V. K., & Sleimen-Malkoun, R. (2015). Effects of task and age on the magnitude and structure of force fluctuations: Insights into underlying neuro-behavioral processes. BMC Neuroscience, 16(1), 1–17. 10.1186/S12868-015-0153-7/FIGURES/6

Wayne, P. M., Gow, B. J., Costa, M. D., Peng, C. K., Lipsitz, L. A., Hausdorff, J. M., Davis, R. B., Walsh, J. N., Lough, M., Novak, V., Yeh, G. Y., Ahn, A. C., Macklin, E. A., & Manor, B. (2014). Complexity-Based Measures Inform Effects of Tai Chi Training on Standing Postural Control: Cross-Sectional and Randomized Trial Studies. PloS One, 9(12). 10.1371/JOURNAL.PONE.0114731

